# Small GTPase patterning: How to stabilise cluster coexistence

**DOI:** 10.1101/477091

**Authors:** Bas Jacobs, Jaap Molenaar, Eva E. Deinum

## Abstract

Many biological processes have to occur at specific locations on the cell membrane. These locations are often specified by the localised activity of small GTPase proteins. Some processes require the formation of a single cluster of active GTPase, also called unipolar polarisation (here “polarisation”), whereas others need multiple coexisting clusters. Moreover, sometimes the pattern of GTPase clusters is dynamically regulated after its formation. This raises the question how the same interacting protein components can produce such a rich variety of naturally occurring patterns. Most currently used models for GTPase-based patterning inherently yield polarisation. Such models may at best yield transient coexistence of at most a few clusters, and hence fail to explain several important biological phenomena. These existing models are all based on mass conservation of total GTPase and some form of direct or indirect positive feedback. Here, we show that either of two biologically plausible modifications can yield stable coexistence: including explicit GTPase turnover, i.e., breaking mass conservation, or negative feedback by activation of an inhibitor like a GAP. Since we start from two different polarising models our findings seem independent of the precise self-activation mechanism. By studying the net GTPase flows among clusters, we provide insight into how these mechanisms operate. Our coexistence models also allow for dynamical regulation of the final pattern, which we illustrate with examples of pollen tube growth and the branching of fungal hyphae. Together, these results provide a better understanding of how cells can tune a single system to generate a wide variety of biologically relevant patterns.

**Author summary:** Where to form a bud? Where to reinforce the cell wall? In which direction to move? These are all important decisions a cell may have to make. Proper patterning of the cell membrane is a critical part of such decisions. These patterns are often specified by the local activity of proteins called small GTPases. Mathematical models have been an important tool in understanding the mechanisms behind small GTPase-based patterning. Most of these models, however, only allow for the formation of a single cluster of active GTPase and thus cannot explain patterns of multiple coexisting GTPase clusters. A previously proposed mechanism for such coexistence can only explain a temporary, unstable coexistence, and fails to explain several key biological phenomena. In this manuscript, we investigate two mechanisms that can produce patterns of many stably coexisting GTPase clusters. Using a combination of modelling techniques, we show why these mechanisms work. We also show that these mechanisms allow for the addition of new clusters to an existing pattern, as is observed for example during the branching of fungal hyphae. With our results, we now have handles to explain the full range of naturally occurring small GTPase patterns.

## Introduction

Many cellular processes must occur at specific locations on the cell membrane. Examples range from the formation of a yeast bud [1], to the localised reinforcements of plant cell walls [2], to coordination of directed cell movement in animals [3]. The localisation of these processes is determined by the local activity of highly conserved small GTPase proteins (e.g., Rho, ROP, Rac, Ras, henceforth referred to as GTPases) [4]. In some cases, such as yeast budding, a single cluster of active GTPase forms, resulting in unipolar polarisation (henceforth referred to as polarisation). In others, e.g., patterned plant cell wall reinforcement, the GTPase pattern consists of many coexisting clusters (Fig 1A). This raises the question how the same biological system can generate different types of patterns. Mathematical models are an important tool in understanding the mechanisms of *de novo* pattern formation, but thus far, most models for GTPase-based patterning only yield polarisation [5, 6].

**Fig 1.**
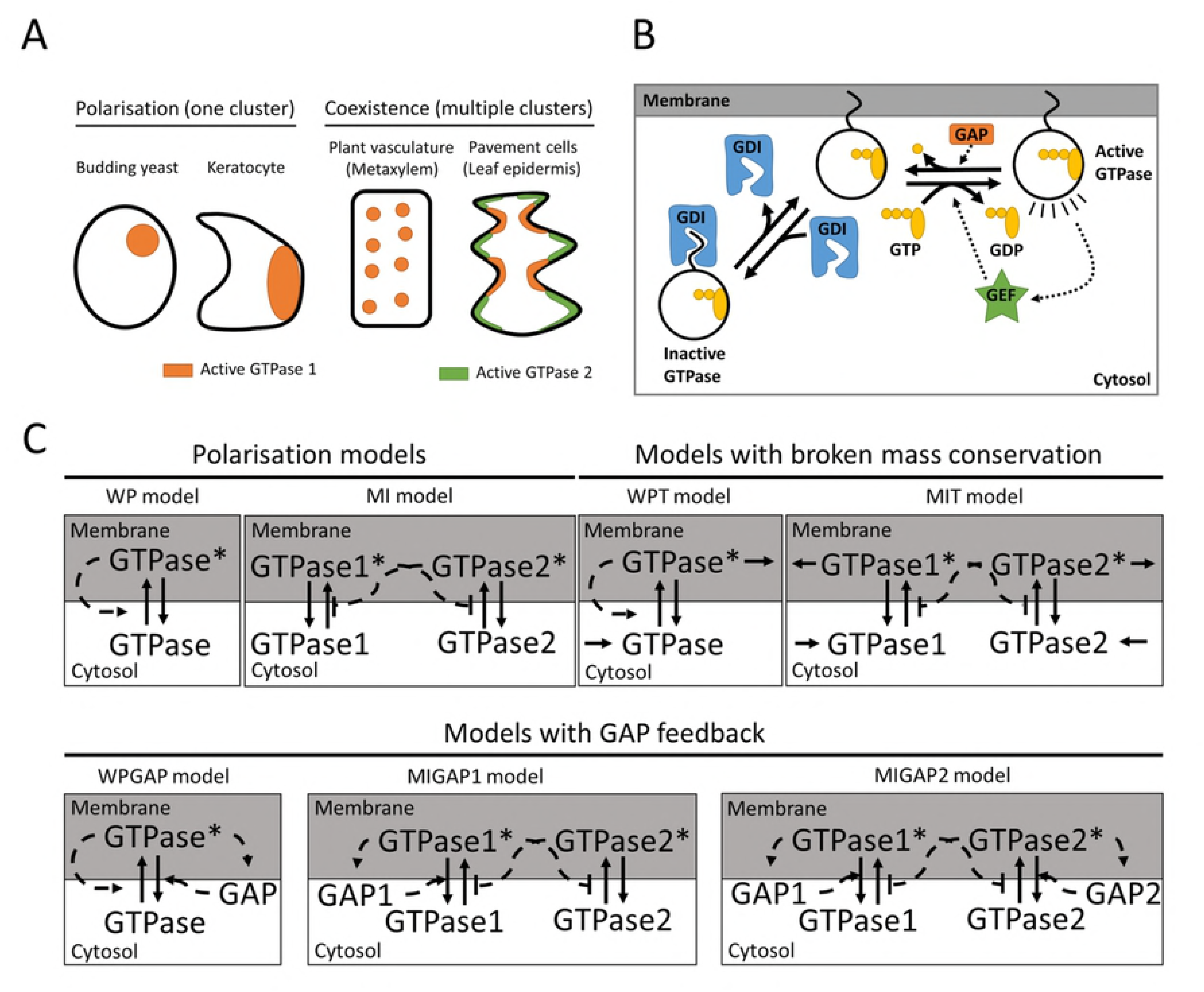
Models for GTPase-based membrane patterning. A: Types of GTPase-based membrane patterns that occur in living cells. Some situations require the formation of a single cluster of GTPase (polarisation, left), whereas others require stable coexistence of multiple clusters (right). B: Active GTP-bound GTPases are inactivated by hydrolysis of GTP to GDP under the influence of GAP proteins. Inactive GTPases can be activated by GEF proteins that promote the exchange of GDP for GTP. GTPases can bind to the cell membrane with their hydrophobic tails, but the inactive form is selectively taken out by GDI proteins. C: Interaction motifs of the reaction diffusion models. Solid lines indicate conversions and dashed lines indicate interactions. Positive interactions are indicated by arrowheads and negative interactions by perpendicular lines. Stars indicate the active form of GTPase.

Small GTPases function as molecular switches with an active, GTP-bound form and an inactive, GDP-bound form (Fig 1B). They can be switched on by Guanine nucleotide Exchange Factors (GEFs), which facilitate exchange of GDP for GTP, and off by GTP hydrolysis, which can be accelerated by GTPase Activating Proteins (GAPs) [7, 8]. The active form is membrane-bound, whereas the inactive form is removed from the membrane by Guanine nucleotide Dissociation Inhibitors (GDIs) [9]. Since diffusion in the cytosol is much faster than at the membrane [10], the inactive form effectively diffuses much faster than the active form. This difference in diffusion is one of the classical ingredients for pattern formation through a Turing-type reaction-diffusion mechanism [11, 12], making these GTPases particularly suitable for membrane patterning in biological systems.

Many theoretical studies have been performed on the role of GTPases in polarisation, for cells of organisms as diverse as animals, yeast, and the cellular slime mold *Dictyostelium discoideum* [5]. Most of these models involve some form of positive feedback, either through direct or indirect self-activation or through double negative feedback between two antagonistic GTPases. In addition, these models generally do not include protein turnover, so GTPase is only interconverted between active and inactive form, and not produced or degraded (generally referred to as *mass conservation*). These two properties seem to form a robust recipe for polarisation [13–17].

However, GTPases also play important roles in the formation of other patterns than simple polarisation. For example, cell wall patterning in at least one common type of plant vasculature, the metaxylem, relies on a pattern of multiple clusters of a GTPase [18–20]. The interdigitated pattern of plant leaf epidermal cells (pavement cells) is under the control of two antagonistic classes of GTPase [21–23]. Furthermore, several GTPases are implicated in the outgrowth of multiple dendrites from neuron cell bodies [4].

Particularly interesting in this context is the role of GTPases in tip growing systems, where the pattern of GTPase clusters may be dynamically regulated, adding an extra layer of complexity. In these systems, a cluster of active GTPase determines the tip region of a growing tube. Some of these systems, such as root hairs [24] and pollen tubes [25], have a single growing tip. In others, such as fungal hyphae, multiple tips can exist simultaneously, each with their own cap of active GTPase. More importantly, new clusters can appear even after others have already been established, resulting in branching. This is important for the establishment of a mycelial network [26, 27]. Additionally, in both pollen tubes and hyphae, growth can proceed in pulses, with the GTPase cap diminishing or disappearing as growth slows or halts [28, 29].

This more complex growth requires the coexistence of multiple clusters as well as the *de novo* generation of new clusters besides existing ones. Studying these processes in one dimension (1D), as is often done for polarisation, saves computational time, but can produce misleading results. For example, many mass conserved reaction-diffusion models for polarisation show phase separating behaviour [13, 30, 31]. Phase separation is characterised by the minimisation of interface length and total curvature in two dimensions (2D), causing clusters to compete until only one remains [32]. In 1D, however, only the (discrete) number of interfaces can be minimised. Consequently, the meta-stable state with multiple clusters that these polarisation models can produce in 1D [13, 30, 31], is likely to be much less stable in 2D.

All this implies that in 2D, multiple domains can only (apparently) stably coexist if supported by an irregular geometry such as the lobe-and-indent shape of leaf epidermal (pavement) cells [33]. A phase separating system, however, cannot explain the initial formation of such a geometry, nor the appearance of additional lobes as these cells grow [34]. These issues of dimensionality and *de novo* cluster formation also apply to a recent theoretical study that proposes that saturation of self-activation, resulting in flat concentration profiles, could slow down the competition between clusters to the extent that they can coexist on biologically relevant time scales [35]. Therefore, a mechanism that allows for truly stable coexistence would offer a more parsimonious explanation for phenomena that require multiple GTPase clusters in a single cell.

From literature we have found two potential ways of obtaining stable coexistence. Firstly, where GTPase-based polarisation models are typically mass conserved, turnover is present in highly similar classical activator substrate-depletion models that do show stable coexistence (e.g., [36]). Mass conservation has been suggested to play an important role in the winner-takes-all mechanism [17] and an adaptation of a polarisation model with production and degradation terms can generate multiple peaks in 1D [37]. These results indicate that sufficiently large GTPase turnover on relevant time scales might explain the stable coexistence of multiple GTPase clusters. Whether that is a reasonable assumption may depend on the system.

Secondly, a parameter regime that allows stable coexistence has been reported without further investigation in a modelling study on negative feedback in polarisation of budding yeast [38]. This suggests a negative feedback that limits the growth of larger clusters may also explain stable coexistence, but this mechanism remains to be fully characterised. Some experimental evidence for negative feedback through the recruitment of GAPs has been found in the case of metaxylem patterning [19].

Here, we aim for a unified understanding of all biologically relevant GTPase-based patterns. Our goal is threefold: (1) to find out which GTPase interaction motifs can make the difference between polarisation and stable coexistence in *de novo* GTPase-based patterning of 2D membranes, (2) to understand why these differences in GTPase interactions lead to different patterns, and (3) to explore the options these mechanisms offer for dynamic regulation of membrane patterning.

We consider the interaction motifs from two existing partial differential equation (PDE) models for polarisation and extensions thereof with GTPase turnover and with negative feedback (See Fig 1C and Table 1 for an overview). These polarisation models are the so-called “Wave Pinning” (WP) model [13] with a single GTPase directly stimulating its own activation, and the “Mutual Inhibition” (MI) model [14] with two GTPases inhibiting each other’s activation. The use of these two different models allows us to draw conclusions that do not depend on the positive feedback mechanism. We break the mass conservation of these models by adding turnover (WPT and MIT, respectively). Negative feedback is included by having mass conserved GTPases activate their own GAP. We consider the WP model with a single GAP (WPGAP) and the MI model both with a GAP for one GTPase (MIGAP1) and GAPs for both GTPases (MIGAP2). To study the entire process of *de novo* cluster formation, we perform simulations starting from a homogeneous state with noise on a 2D domain sufficiently large for the formation of complex patterns. We found that both polarisation models indeed robustly yield a single GTPase cluster, whereas either breaking mass conservation or adding GAP feedback appears to be sufficient for stable coexistence. We also provide insight into the way these mechanisms operate, by using ordinary differential equation (ODE) models derived from the PDEs to study fluxes between competing clusters. Finally, we revisit the tip growing systems to explore the options that the different mechanisms offer for dynamically regulating established patterns. In particular, our findings reveal conditions that allow for branching.

**Table 1.** Characteristics of the different models. WP = wave pinning, MI = mutual inhibition, WPT = WP with turnover, MIT = MI with turnover, WPGAP = WP with GAP feedback, MIGAP1 = MI with GAP feedback on one GTPase, MIGAP2 = MI with GAP feedback on both GTPases.

| Model | Positive feedback | Mass conservation | Number of GTPases | Number of GAPs | Number of variables |
| --- | --- | --- | --- | --- | --- |
| WP | Direct | Yes | 1 | 0 | 2 |
| MI | Double negative | Yes | 2 | 0 | 4 |
| WPT | Direct | No | 1 | 0 | 2 |
| MIT | Double negative | No | 2 | 0 | 4 |
| WPGAP | Direct | Yes | 1 | 1 | 4 |
| MIGAP1 | Double negative | Yes | 2 | 1 | 6 |
| MIGAP2 | Double negative | Yes | 2 | 2 | 8 |

## Results

### Simple mass conserved models can only result in polarisation

In this section we study the wave pinning (WP) and mutual inhibition (MI) models, which have in common that the total amount of GTPase is conserved. Both can be written in the following dimensionless form (see section 1 of S1 Appendix):

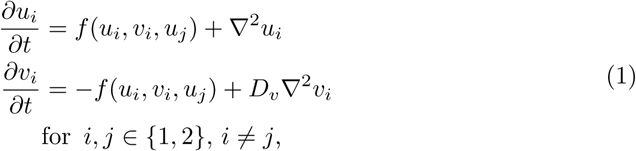

where *u_i_* and *v*_*i*_ are the concentrations of the *i*^th^ active and inactive GTPase respectively, *D*_*v*_ is the ratio between diffusion coefficients of inactive and active GTPase (*»* 1), and *f* represents the interconversion between active and inactive form. For the WP model *u*_*i*_ = *u*_*j*_ = *u* and *v*_*i*_ = *v*_*j*_ = *v*, and function *f* consists of constant activation and inactivation terms and a saturating self-activation term. The dimensionless form of *f* is given by:

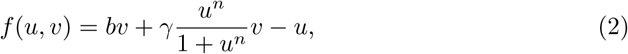

where *b* is a constant activation rate, *γ* the self-activation rate at saturation, and *n* the hill coefficient describing saturating self-activation. Due to mass conservation, the average total (dimensionless) GTPase concentration *T* is a constant determined by initial conditions.

For the MI model, function *f* has constant activation and inactivation terms and a saturating inhibition term. The dimensionless form is given by:

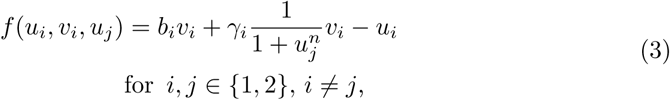

where *b*_*i*_ is the constant activation rate of GTPase *i*, *γ*_*i*_ the activation rate of GTPase *i* that can be inhibited by GTPase *j*, and *n* the hill coefficient describing inhibition. Again, the total amount of each GTPase is constant and we use the average total GTPase concentrations *T*_1_ and *T*_2_ as parameters.

We simulated these models on a rectangular domain with periodic boundary conditions in the horizontal direction and zero-flux boundary conditions in the vertical direction. In this way, our domain is an open cylinder resembling the membrane of a large, elongated cell. Domain sizes were large compared to those used by the original WP model study on polarisation [13], but the other parameters had similar values (Table 2). We enlarged the domain size to ensure domain size was not limiting the formation of multiple clusters. All simulations started in the homogeneous steady state (HSS) with a small amount of noise added (see Methods for details).

**Table 2.** Descriptions and default values of parameters used for the dimensionless PDE models. In case of multiple GTPases or GAPs the same values for *b*, *γ*, *c*, *d*, *ξ*, and *T*_*g*_ were used for both variants. N.A. = not applicable.

| Parameter | Description | Model |  |  |  |  |  |  |
| --- | --- | --- | --- | --- | --- | --- | --- | --- |
|  |  | WP | MI | WPT | MIT | WPGAP | MIGAP1 | MIGAP2 |
| $b$ | Base activation rate | 0.1 | 0.1 | 0.1 | 0.1 | 0.1 | 0.1 | 0.1 |
| $\gamma$ | Feedback activation rate | 2 | 2 | 2 | 2 | 2 | 2 | 2 |
| $n$ | Hill exponent of feedback saturation | 2 | 2 | 2 | 2 | 2 | 2 | 2 |
| $c$ | GAP activation rate | N.A. | N.A. | N.A. | N.A. | 1 | 1 | 1 |
| $d$ | GAP inactivation rate | N.A. | N.A. | N.A. | N.A. | 1 | 1 | 1 |
| $T_{(1)}$ | Average concentration of total GTPase (1) | Variable | | | | | | |
| $T_2$ | Average concentration of total GTPase 2 | N.A. | 5 | N.A. | N.A. | N.A. | 5 | 20 |
| $\sigma_{(1)}$ | Production rate of GTPase (1) | Variable | | | | | | |
| $\sigma_2$ | Production rate of GTPase 2 | N.A. | N.A. | N.A. | 0.2 or 0.4 | N.A. | N.A. | N.A. |
| $\xi$ | GTPase degradation rate | N.A. | N.A. | 0.1 | 0.1 | N.A. | N.A. | N.A. |
| $T_g$ | Average concentration of total GAP | N.A. | N.A. | N.A. | N.A. | 10 | 10 | 10 |
| $D_v$ | Ratio diffusion coefficients of inactive to active GTPase | 100 | 100 | 100 | 100 | 100 | 100 | 100 |
| $D_G$ | Ratio diffusion coefficients of active GAP to active GTPase | 100 | 100 | 100 | 100 | 100 or 40 | 100 | 100 |
| $D_g$ | Ratio diffusion coefficients of inactive GAP to active GTPase | 100 | 100 | 100 | 100 | 100 | 100 | 100 |

As previously predicted [13, 17], both polarisation models consistently generate a single cluster of active GTPase in a pattern that changes from a spot to a stripe to a gap as the amount of GTPase increases (Fig 2). Transiently, multiple clusters of varying sizes form in an irregular pattern, with the larger clusters growing at the expense of the smaller ones (Fig 3, S1 Video). Because of the varying sizes, most clusters disappear quickly and only the last few compete for a long time. This shows that the previously proposed mechanism of slow competition [35] cannot explain *de novo* formation of patterns of many coexisting clusters.

**Fig 2.**
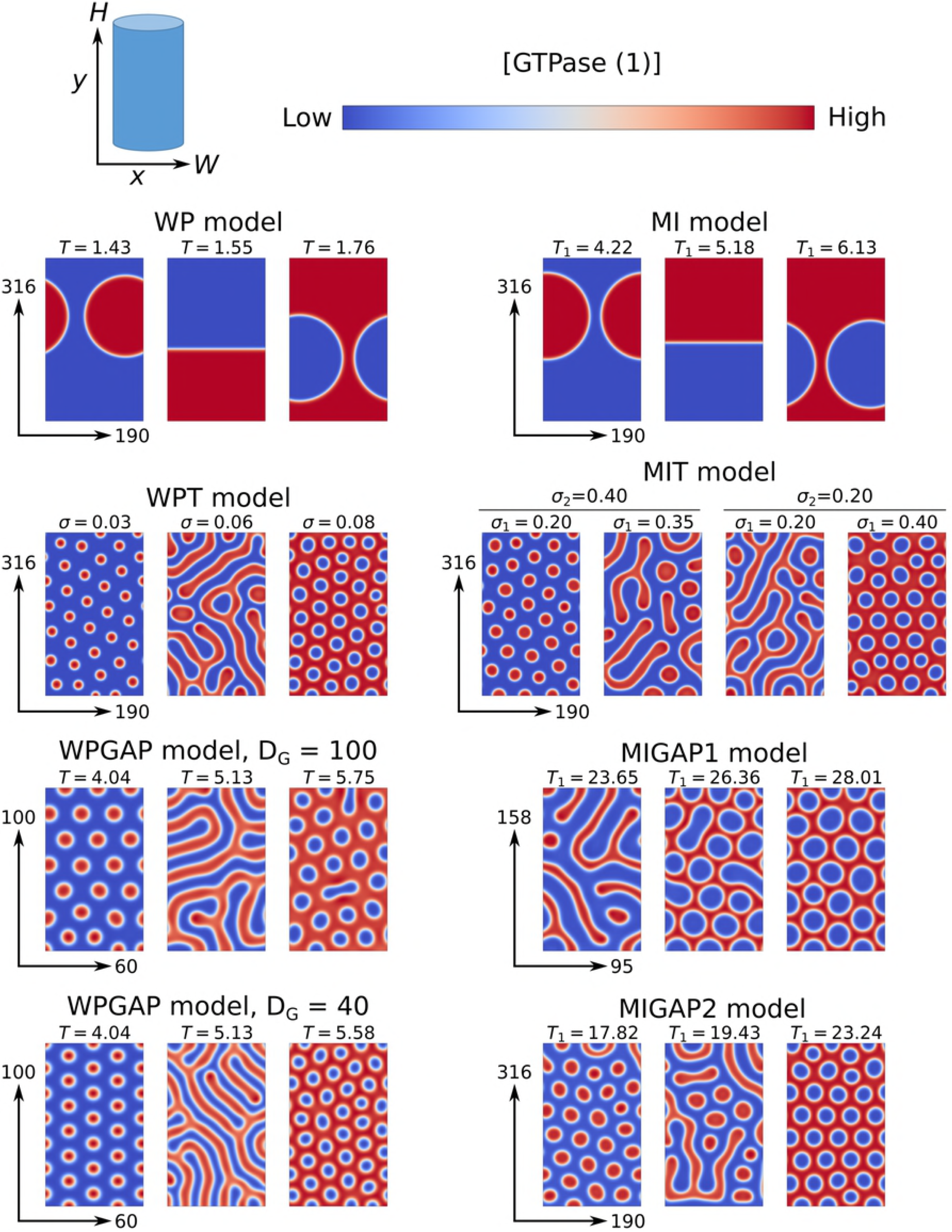
Steady state active GTPase concentrations ([GTPase]) obtained by model simulations. *T* = total amount of GTPase, *σ* = GTPase production rate. All simulation domains have periodic boundary conditions in the x-direction and zero flux boundary conditions in the y-direction. Dimensionless domain heights (*H*) and widths (*W*) are indicated by arrows. For models with two GTPases, only the concentrations of the first GTPase are shown. In these cases, the second GTPase always clusters where the first does not. Default parameters (Table 2) were used with indicated values of *T* and *σ*. For the WPGAP model, two different values were used for the active GAP diffusion coefficient (*D*_*G*_). Concentration ranges and simulation end times for each simulation are shown in S1 Fig.

**Fig 3.**
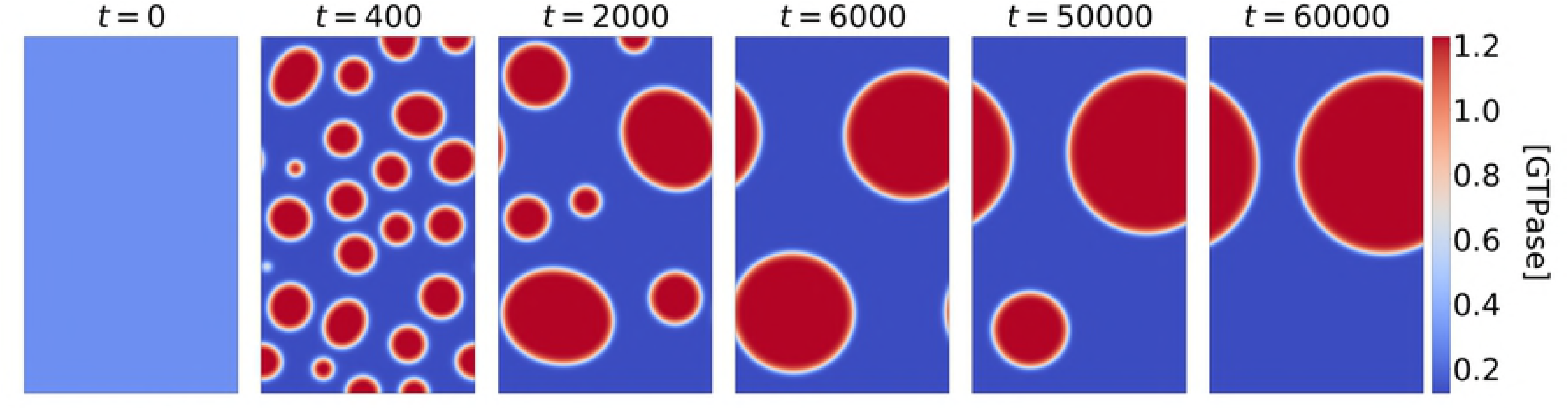
Transient active GTPase concentrations ([GTPase]) from simulations of the WP model with. *T* = 1.43. Simulation conditions were as in Fig 2.

Linear stability analysis (LSA, see Methods) performed on the WP model, reveals that the domain size used does not prevent multiple different wave numbers from becoming unstable (S2 Fig). Therefore, the formation of a single cluster results from dynamics that occur after the initial symmetry breaking, when linearisation is no longer appropriate. This means, as previously noted [39], that LSA cannot be used to determine the length scale of the final pattern, because it is only valid around the homogeneous state.

To visualise regions of parameter space where the homogeneous state is unstable and patterns form spontaneously (so-called Turing regimes), we generated two parameter bifurcation plots. For models with two parameters this can be done with LSA. For larger models this is no longer feasible and therefore we also used the more approximate local perturbation analysis (LPA, see Methods). Where both methods can be used, they yield highly similar results (Fig 4). For the WP model, they reveal that patterns only form if the self-activation parameter *γ* is sufficiently large, while the total amount of GTPase must be within certain boundaries. For the MI model, LPA predicts that a wide range of total GTPase amounts give rise to spontaneous pattern formation, as long as the amounts of both GTPases are similar (Fig 5A).

**Fig 4.**
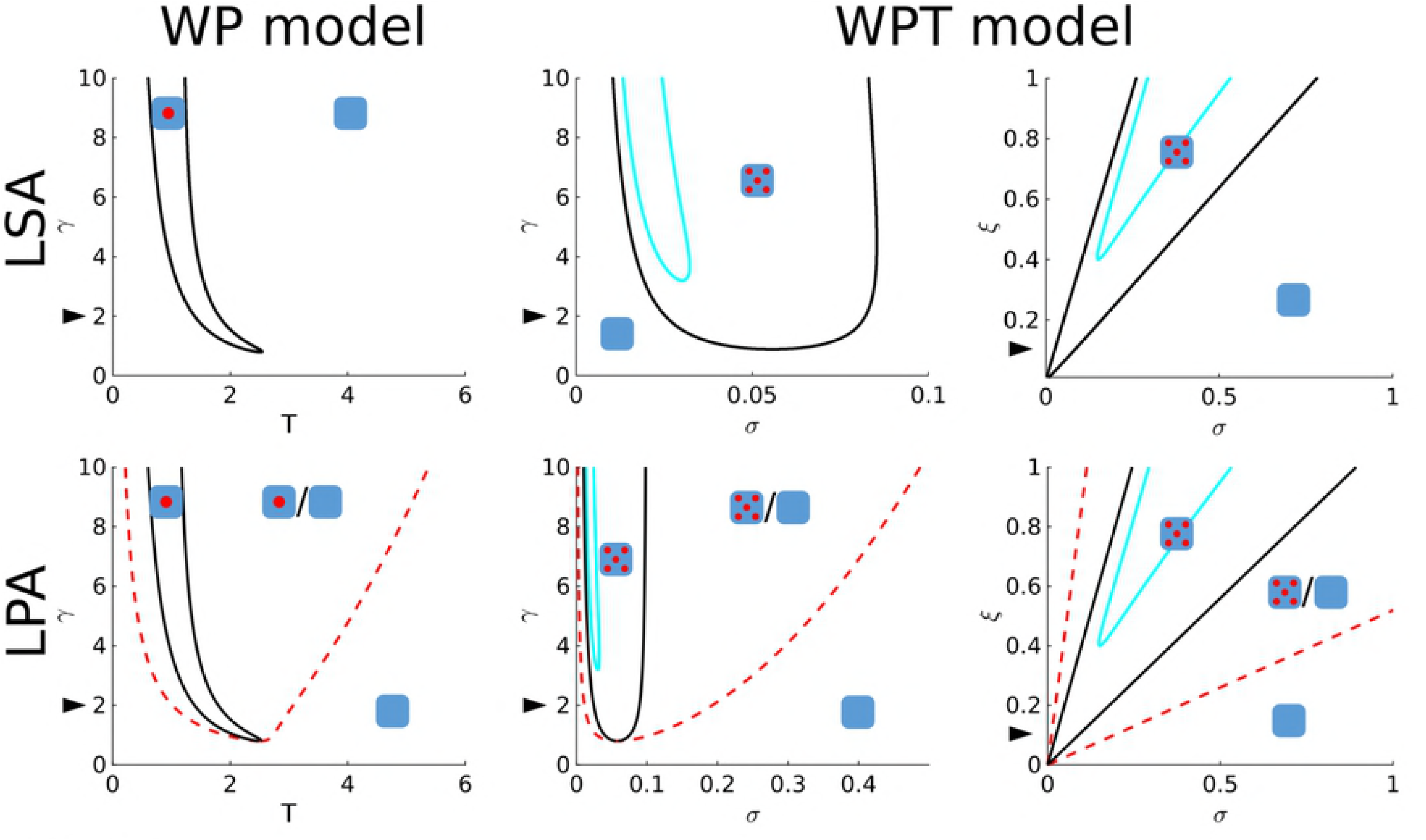
Two parameter bifurcation plots for models with two variables (WP and WPT). Both LSA (top) and LPA (bottom) analyses were performed on the WP model for parameters *T* and *γ* (left) and on the WPT model for parameters *σ* and *γ*, and *σ* and *ξ* (right). Cartoons indicate kind of pattern formed in different regimes (empty square: no pattern, single spot: polarisation, five spots: coexistence). Black lines delimit Turing regimes in which the homogeneous steady state is unstable. Dashed red lines (LPA only) delimit predictions of regimes in which heterogeneous states exist that may be reached from the homogeneous state by an arbitrarily large local perturbation. Cyan lines indicate Hopf regimes. Arrowheads indicate default parameters.

**Fig 5.**
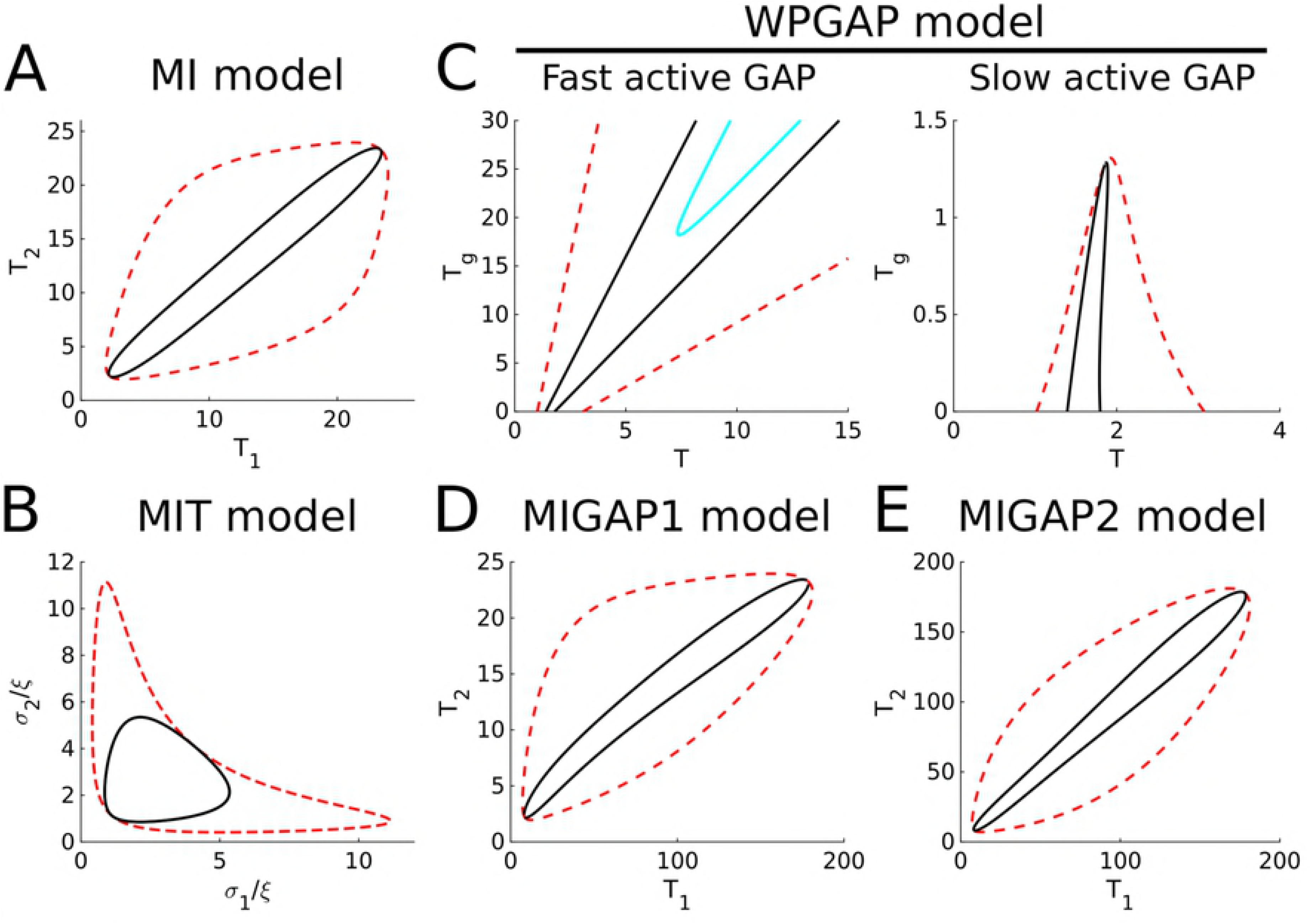
Two parameter bifurcation plots for LPA analyses on models with more than two variables. Default parameters were used. The bifurcation plot for the two ratios *σ/ξ* in the MIT model (B) appears to be the same regardless of the value of *ξ*_1_ = *ξ*_2_ = *ξ* (tested for *ξ ∈* {0.05, 0.1, 0.2, 1}). For the WPGAP (C) model, the active GAP diffusion coefficient (*D*_*G*_) was considered both in the limit *D_G_* → ∞ (Fast active GAP) and in the limit *D*_*G*_ → 0 (slow active GAP). Lines delimiting different regimes are coloured as in Fig 4. For the MI models with GAP feedback (D and E) we only considered fast active GAP.

### Breaking mass conservation is sufficient to allow stable coexistence of multiple active GTPase clusters

To investigate the effect of breaking mass conservation, we extended the two polarisation models with production and degradation terms. Since translation is a cytosolic process, we assume that GTPases are produced in their inactive form. Following a previous study [37], we here consider degradation of the active form. Later, we will also consider degradation of the inactive form. These assumptions result in the following dimensionless model equations:

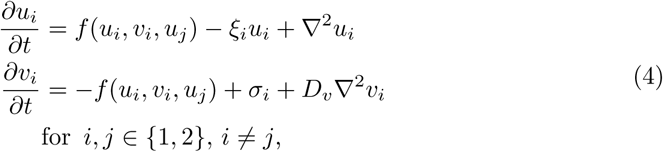

with constant production *σ*_*i*_ of the inactive form, degradation of the active form with factor *ξ*_*i*_, and *f* given by Eq 2 or Eq 3. The equations for the WPT model are equivalent to those used in a previous study [37].

Consistent with previous results in 1D [37], breaking mass conservation allows multiple clusters to survive (Fig 2, S1 Video). As for classical Turing systems [40], higher GTPase production shifts the pattern from spots to stripes to gaps, suggesting that the most important difference between these models and polarisation models is indeed mass conservation. In the limit of no turnover, the WPT model converges to the WP model. Simulations show a gradual decrease in the number of stably coexisting clusters per unit area with decreasing turnover, until a single one remains and polarisation behaviour is recovered (S3 Fig).

As for the WP model, LSA performed on the WPT model reveals many different unstable wave numbers (S2 Fig), although simulations show a different class of patterns. Bifurcation analysis reveals that, for patterning to occur, the WPT model also requires a sufficiently large self-activation parameter *γ*. In addition, the GTPase production rate *σ* must be within certain boundaries determined by degradation rate *ξ* (Fig 4). Bifurcation analysis also reveals a Hopf regime within the Turing regime for the WPT model. Simulations in the Hopf regime reveal no oscillations of the final pattern (S2 Video), likely because this regime only applies close to the homogeneous state. LPA shows that the Turing regime for the MIT model is considerably less elongated than that for the MI model (Fig 5B). In addition, the Turing regime appears to scale linearly with the degradation rate.

Both models considered here assume degradation of the active form. Similar bifurcation and simulation analyses on a model with degradation of the inactive form do not reveal regimes that admit stable patterns (see section 5 in S1 Appendix). This shows that turnover can only stabilise multiple coexisting clusters if the active form is (also) degraded. Such degradation could occur, for example, by recycling of membrane patches.

### Adding GAP feedback to the mass conserved models is also sufficient to allow for coexistence

To investigate the effect of negative feedback, we modified both the WP and MI model by including GAP proteins in such a way that both the total amount of GTPase and the total amount of GAP are conserved. For the MI model, we considered both cases of a single GAP acting on one of the GTPases (MIGAP1) and one GAP for each GTPase (MIGAP2). This results in the following dimensionless equations for the GAP models:

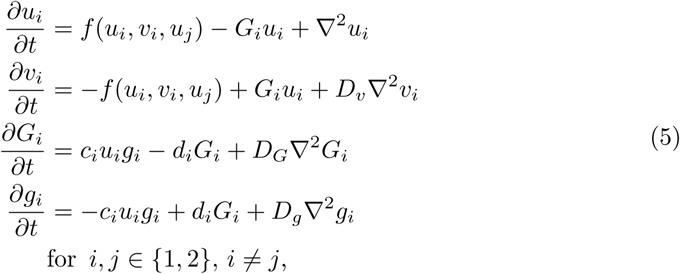

where *G_i_* and *g*_*i*_ are the dimensionless concentrations of active and inactive GAP respectively, *D*_*G*_ and *D*_*g*_ are the diffusion coefficients of active and inactive GAP relative to that of active GTPase respectively, *c*_*i*_ is the GTPase-dependent GAP activation rate, and *d*_*i*_ is a constant GAP inactivation rate. Function *f* is again defined by Eq 2 or Eq 3. For the MIGAP1 model, the second GAP (*G*_2_ and *g*_2_) is absent. Like the total amount of GTPase, the total amount of GAP is conserved, resulting in additional parameters for the average total GAP concentration *T*_*g,i*_.

Consistent with previous indications [38], addition of GAP feedback results in similar patterns as found for breaking mass conservation: multiple clusters of active GTPase become stable and the pattern shifts from spots to stripes to gaps for increasing levels of total GTPase (Fig 2, S1 Video). For the MI model, adding GAP feedback to only one of the two GTPases is sufficient to achieve this effect. The similar response of the two different polarisation models indicates that the difference between polarisation and coexistence does not depend on the self-activation mechanism.

Bifurcation analysis of the WPGAP model reveals that, as long as GAPs diffuse fast compared to active GTPase, the Turing regime widens and shifts to higher amounts of GTPase as the amount of GAP increases (Fig 5C). This is consistent with previous indications that negative feedback may act as a buffer against fluctuations in GTPase concentration [38]. However, when active GAP diffusion is equal to that of active GTPase, the picture changes completely and GAPs severely hamper pattern formation. Trial simulations in the narrow Turing regime all resulted in polarisation (S4 Fig). Therefore, it seems that for GAP feedback to stabilise coexistence, the effective diffusion constant of active GAP must be larger than that of active GTPase. However, formation of coexisting GTPase clusters is still possible if active GAP diffuses more slowly than inactive GTPase (Fig 2). A Hopf regime can also be found for the WPGAP model. As for the WPT model, simulations in the Hopf regime do not reveal any oscillations of the final pattern (S2 Video).

For the MIGAP2 model, there is a wide range of total GTPase amounts for which patterns form spontaneously, as long as the total amount of both GTPases is similar (Fig 5E). For the MIGAP1 model, a similar range is present, but there must be considerably more of the first GTPase than of the second, because the second is not hindered by GAP feedback (Fig 5D).

### Multiple clusters drawing GTPase from a homogeneous pool compete until only the largest survives

To better understand why some models yield only polarisation, whereas others allow for multiple stable clusters, we considered a simplified model describing the competition between multiple clusters, as transiently generated by the polarisation models (Fig 3, S1 Video). This approximation treats each cluster as a compartment, with an additional non-cluster compartment representing a global pool of inactive GTPase. By using compartments, we implicitly assume that the clusters have sharp boundaries and constant active GTPase concentrations, making the total amount of GTPase in a cluster proportional to its area. Using these assumptions, we derived a system of ODEs from the PDE models (for derivation see section 6.1 of S1 Appendix), which is comparable to a more phenomenological two-cluster model presented by Howell et al. [41]. Clusters recruit active GTPase from the inactive pool at a rate proportional to their area or, equivalently, their total amount of GTPase (Fig 6A). They lose GTPase both at a constant rate, reflecting inactivation, and at a rate proportional to the circumference of the cluster, reflecting loss at the boundary (Fig 6A).

**Fig 6.**
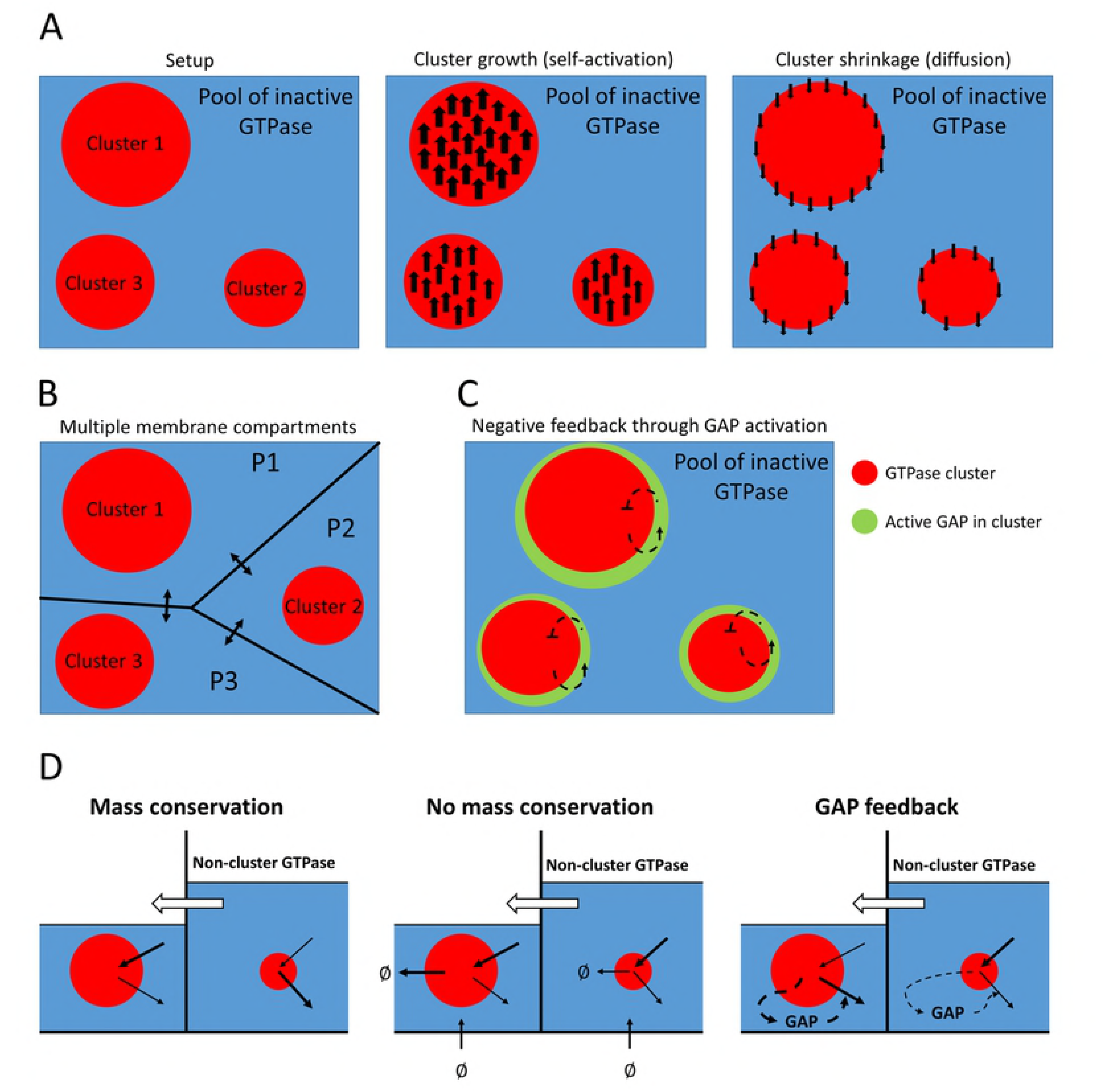
Principles of the simplified ODE models. A: The basic mass conserved ODE model starts with an arbitrary number of differently sized clusters (left, three are shown). Clusters compete for the inactive GTPase from a shared homogeneous pool. Activation and inactivation of membrane GTPase occur proportional to cluster size (middle). Additional loss of active GTPase (by diffusion from cluster to pool) occurs proportional to cluster circumference (right). B: An adaptation of the basic ODE model where each cluster has its own local pool of inactive GTPase (P1-P3) with fluxes connecting these local compartments (arrows). C: An adaptation of the basic ODE model with GAP feedback, where each cluster recruits GAP from a shared homogeneous pool at a rate proportional to the cluster size and GAPs (green) inactivate the GTPase in the cluster. D: Larger clusters more effectively deplete their local pool (blue), resulting in a concentration gradient and corresponding flux of GTPase from compartments with small clusters to compartments with large clusters. With mass conservation, this process continues until the small cluster is depleted. GTPase turnover effectively redistributes GTPases by removing more from larger clusters and providing new GTPase homogeneously, allowing the smallest cluster to compete in spite of this flux. At the same time, the larger cluster suffers from a smaller GTPase pool to recruit from. When GAP feedback is added, larger clusters recruit more GAP enhancing their own depletion, allowing smaller clusters to compete.

The resulting basic ODE model is given by:

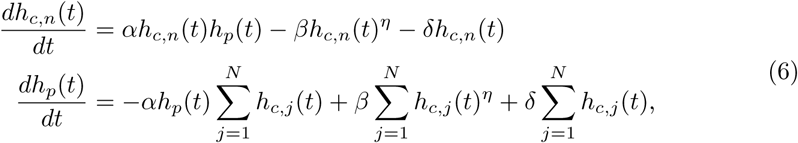

where *h*_*c,n*_ is the amount of (active) GTPase in cluster *n* (which is proportional to the area of cluster *n*), *h*_*p*_ is the amount of GTPase in the inactive pool, *N* is the total number of clusters, exponent *η* is 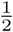 for circular clusters, *α* is a positive constant determining self-activation, *β* is the rate at which clusters lose GTPase by diffusion across the circumference, and *δ* is a constant inactivation rate. This system is redundant, because the total amount of GTPase is conserved 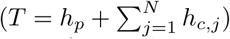.

We calculated the rate of change of the ratio between the sizes of two arbitrary clusters *i* and *k*:

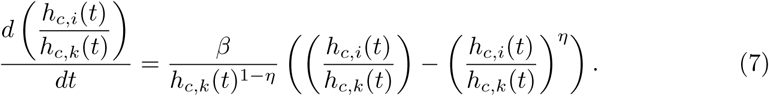

Not only for 2D circular clusters 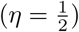, but for any 0 *< η <* 1, the ratio *h_c,i_/h*_*c,k*_ always increases when *h_c,i_ > h*_*c,k*_ and always decreases when *h_c,i_ < h*_*c,k*_, so that differences in cluster size will always grow. This suggests that a single winner will emerge, and this winner will be the cluster that started out as the largest. Numerical simulations confirm this, even when the initial difference in cluster size is very small (Fig 7A). These results show that polarisation will occur when cluster growth increases with cluster size faster than shrinkage does and all clusters rely on the same homogeneous pool of inactive GTPase.

**Fig 7.**
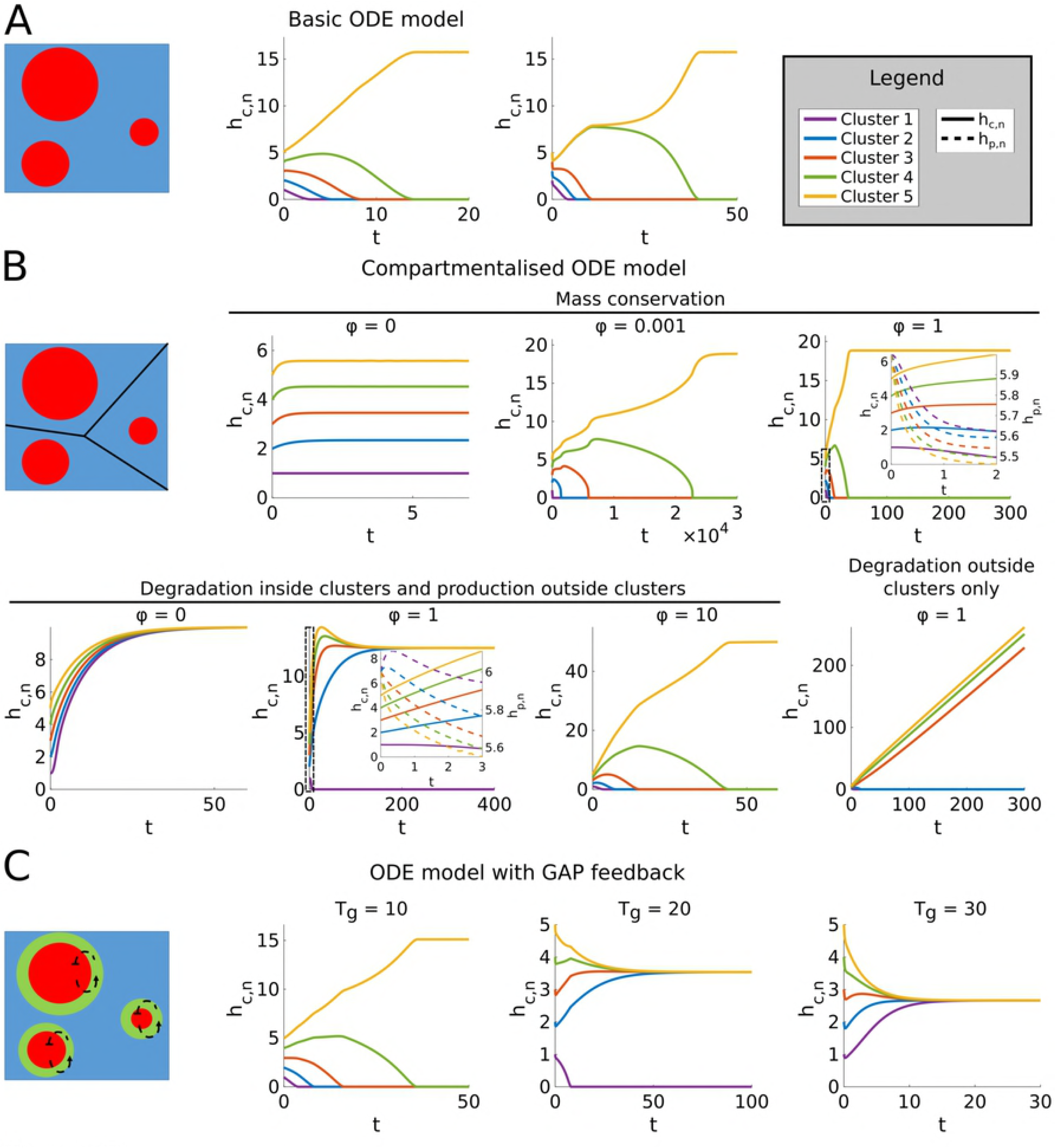
Simulations of the ODE models. Cartoons correspond to model variants described in Fig 6. All simulations started with five clusters of different size (*h*_*c,n*_, *n* ∈ {1, 2, 3, 4, 5}). Parameters *α*, *β*, and *ε* can be scaled out and were set to 1. Other parameter values were: *δ* = 5, *η* = 0.5, and *γ* = *ζ* = 10. A: Simulations of the basic mass conserved ODE model. The total amount of GTPase in the system (*T*) was set to 21. B: Simulations of the ODE model with membrane compartments. Clusters had starting levels of GTPase of 1, 2, 3, 4, and 5, while all membrane compartments had a starting level of 6. Exchange rates between compartments (*φ*) were varied for both mass conserved and non-mass conserved cases. For cases without mass conservation, GTPase production rates in the local pools were set to 1 and degradation rates in either clusters or local pool were set to 0.1. Insets show details of dynamics in boxed regions, including dynamics of GTPase in local pool compartments (*h*_*p,n*_, dashed lines). C: For GAP simulations, total GAP (*T*_*g*_) was varied, always starting with all GAP in the shared pool.

### With GTPase turnover, smaller clusters can sustain themselves from their own local supply

The previous result does not depend on mass conservation, since, if present, the terms describing production and degradation cancel out in the derivation of Eq 7. To understand the mechanism by which breaking mass conservation stabilises coexistence, we therefore have to take into account that the competing clusters actually are spatially separated, possibly resulting in local differences in inactive GTPase availability. We therefore extended the basic ODE model in Eq 6 by giving every cluster its own local pool with an amount *h*_*p,n*_ of inactive GTPase (Fig 6B, see section 6.2 of S1 Appendix for details). Clusters can only draw GTPase from their own compartments. Inactive GTPase is passively exchanged between compartments at a constant rate resembling Fick’s law for diffusion. This results in:

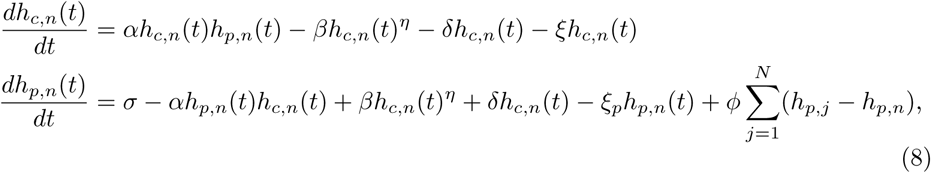

where *σ* is a constant production rate, *ξ* and *ξ*_*p*_ are constant degradation rates of active and inactive GTPase respectively, and *φ* is the constant exchange rate between compartments due to diffusion of inactive GTPase.

The counterpart of Eq 7 now reads as:

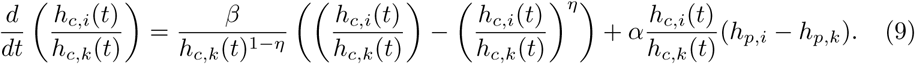

The extra term depends on the difference between the amounts of GTPase in the two non-cluster compartments. Since larger clusters are expected to more effectively drain their local compartment, this term will act to decrease the ratio between sizes of clusters *i* and *k* if cluster *i* is larger and decrease this ratio if cluster *k* is larger. However, it is not a priori clear which of the two terms in Eq 9 is dominant. This dominance may be affected by turnover.

Simulations suggest that without production and degradation (*σ* = *ξ* = *ξ*_*p*_ = 0, mass conservation) the compartmentalised model only allows coexistence in the trivial case where the exchange rate between compartments is zero (Fig 7B). In all other cases, competition proceeds until only a single cluster remains, even though inactive GTPase levels outside large clusters are smaller than GTPase levels outside small clusters. These results indicate that in the mass conserved case larger clusters do indeed more effectively deplete their local GTPase pool. Although this hampers their further growth, it also results in a gradient and corresponding net flux of non-cluster GTPase from compartments with smaller clusters to those with larger clusters (Fig 6D). Since there is nothing to disrupt this flux or replenish the GTPase pools for smaller clusters, this process continues until only the largest cluster remains.

Simulations with turnover, but without degradation of non-cluster GTPase did yield coexistence (Fig 7B). Therefore, disruption of the flux form smaller to larger clusters by turnover does not seem to be required for coexistence. Instead, turnover seems to stabilise coexistence by removing GTPase mostly from larger clusters and redistributing it homogeneously, thereby compensating the diffusive flux to the domains of larger clusters (Fig 6D). At the same time, the growth of the larger clusters is hampered by their smaller pool of non-cluster GTPase, allowing the smaller clusters to catch up. The coexistence was lost at high rates of exchange between compartments, where the system converges to the basic model with a single homogeneous pool.

Simulations with degradation in non-cluster compartments instead of in clusters all resulted in unbounded growth of surviving clusters, again suggesting that removal must target active GTPase to allow stable patterns to form. Note that this unbounded growth is an artefact of the simplified model. In reality, described by the full spatial (PDE) model, cluster growth stops when all clusters have merged, forming a new homogeneous state (see section 5 in S1 Appendix and S3 Video). When considering degradation of both active and inactive form at the same time, stable coexistence can be found (section 7 in S1 Appendix and S4 Video), indicating that degradation of the inactive form does not preclude coexistence.

### GAP feedback stabilises coexistence by punishing larger clusters

To study the way in which GAPs stabilise coexistence of multiple GTPase clusters, we also considered an extension of the basic ODE model from Eq 6 including the effect of GAPs (Fig 6C). In this extension, every cluster has its own amount of active GAP *G*_*c,n*_ and all clusters share a common pool with an amount *G*_*p*_ of inactive GAP. Active GAPs inactivate GTPase at a rate proportional to GAP and GTPase concentrations. GAPs are activated by the active GTPase clusters, and inactivated at a constant rate and by diffusion across the boundaries of the cluster. Together, these assumptions lead to the following extended model:

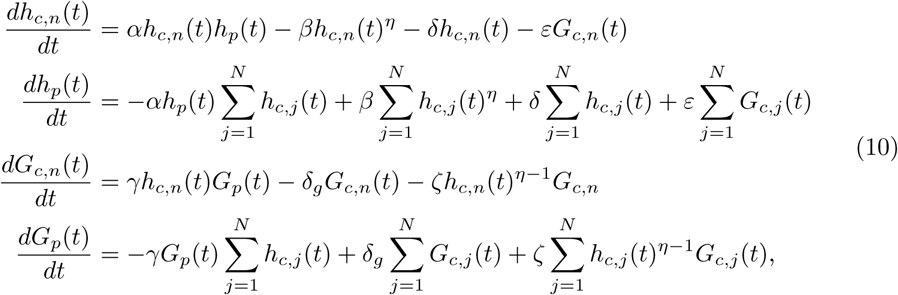

where *G*_*c,n*_ is the total amount of active GAP in cluster *n*, *γ* is the GTPase-dependent GAP activation rate, *δ*_*g*_ is a constant GAP inactivation rate, *ζ* is the rate at which the cluster loses GAP by diffusion across the circumference, and *ε* the rate of GAP-dependent GTPase inactivation. The form of the GAP-related terms in these equations is a direct consequence of using amounts instead of concentrations (see section 6.3 of S1 Appendix). The total amount of GAP 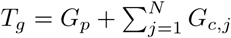 is conserved.

In the same way as before, we obtained for each pair (*i, k*) of clusters:

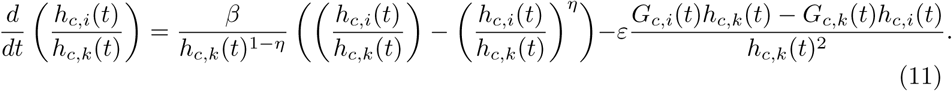

Due to the extra GAP-dependent term, differences in cluster size no longer always increase. As the cluster size increases, the amount of GAP in the cluster will also increase, changing the rate at which the ratio between the two cluster sizes changes in favour of the smaller cluster. However, the net effect of the GAP-dependent term depends on the product of the amount of GAP in one cluster and the size of the other, so the sign of this term is not a priori clear. If we assume that GAP dynamics is fast compared to changes in cluster size, we can take a quasi steady state approximation, which allows the amounts of GAP to be written as a function of cluster size (see section 8 of S1 Appendix). This way Eq 11 can be written as:

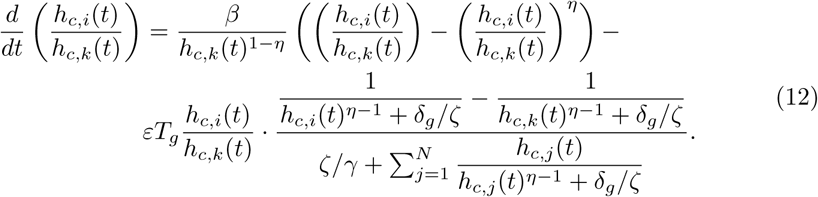

Here, the first term always acts to increase differences in cluster size, whereas the second term always acts to decrease differences for the entire range of 0 *< η <* 1. Which effect dominates depends on the parameters. The equation suggests that a larger total amount of GAP will shift the balance in favour of coexistence. Numerical simulations confirmed this (Fig 7C). This result indicates that GAP feedback stabilises the coexistence of multiple clusters by punishing larger clusters, which tend to activate more GAPs (Fig 6D).

Since the GAP dependent term in Eq 12 is proportional to the total amount of GAP, one might expect that the number of stably coexisting clusters can be increased by simply increasing the amount of GAP. Numerical simulations of the ODE model seem to confirm this (Fig 7C). However, the ODE model assumes the existence of clusters which in the full PDE model is not guaranteed. This provides the additional constraint that the parameters must remain in the Turing regime, which strongly limits the extent to which the total amount of GAP can be increased (Fig 5C).

### Dynamic regulation of established patterns: a case study of tip growing systems

Above, we established two mechanisms that can lead to stable coexistence of multiple GTPase clusters. As a case study, we now explore tip growing systems, where dynamical regulation of the GTPase pattern is often important. In pollen tube tip growth, the supply of GAP increases as the cap of active GTPase at the tip grows, so that the size of the GTPase cluster oscillates and the tip grows in pulses [28, 42]. In the fungus *Ashbya gossypii*, the tip growth complex (polarisome) sometimes disappears, corresponding with a stop in tip growth, after which it spontaneously re-establishes and growth continues, suggesting a negative feedback [29]. In addition, two types of branching occur in *A. gossypii*: lateral branching, where a new tip appears somewhere along the length of an existing hypha, and apical branching, where a growing tip splits in two [43]. We use proof of principle simulations of single GTPase clusters both to explore the options our mechanisms give for such dynamic regulation of the GTPase pattern and to offer possible explanations of these phenomena (see Methods for implementation details).

These simulations show that the cases of pulsing and disappearing GTPase caps can be reproduced by an increase in either the total amount of GAP (Fig 8A-E, S5 Video), or the GTPase degradation rate (S5 FigA-E, S5 Video). If these parameters return to their base levels after the cluster has shrunk or disappeared, it will immediately grow back or reappear, allowing the cycle to start again. The required change in parameters under the current settings is at most 50%, which could reasonably be achieved by changes in GAP production or release, or the recycling of membrane proteins. Note that as long as we start off with an existing cluster, the total amount of GAP or the GTPase degradation rate may even end up somewhat outside the Turing regime without the cluster disappearing. However, the cluster disappears long before leaving the regime where LPA predicts a heterogeneous state to exist (Fig 8B and S5 FigB). This indicates that although Turing regimes are accurately predicted by LPA, regimes with a stable heterogeneous state are not.

**Fig 8.**
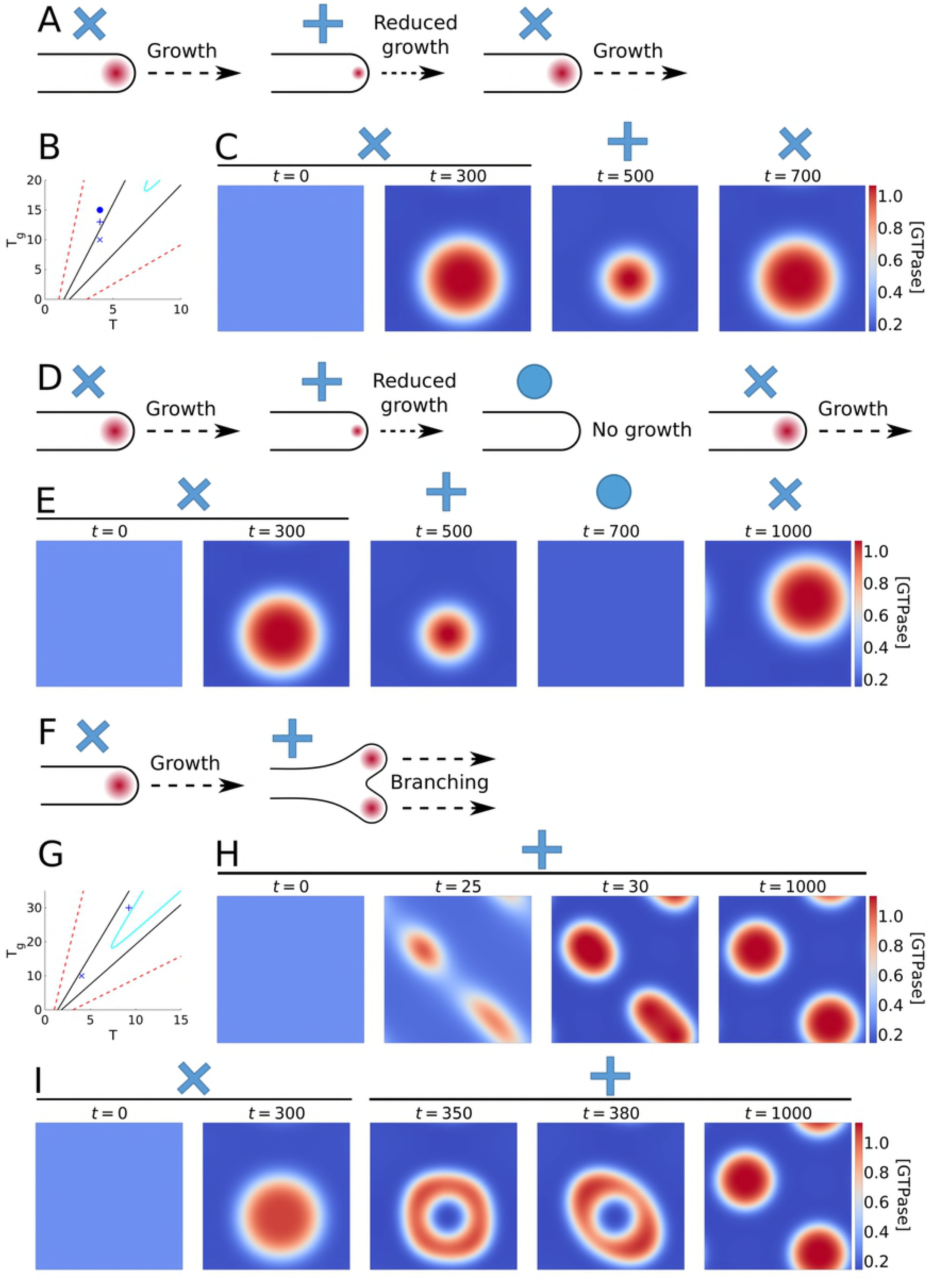
Tip growth simulations with the WPGAP model. B and G: Bifurcation plots with symbols showing parameter settings used in the simulations, corresponding to symbols above simulations and cartoons. For meaning of lines see Fig 5. A: In pollen tubes, growth occurs in pulses as negative feedback during growth results in an oscillatory GTPase cluster at the growing tip. C: Simulation of a single active GTPase cluster with increase in *T*_*g*_ upon reaching steady state followed by a return to starting levels. D: In growing hyphae of *A. gossypii*, the tip growth complex sometimes disappears corresponding to a halt in growth, suggesting involvement of negative feedback. E: Simulation with two subsequent increases in *T*_*g*_, followed by a return to the starting level. F: In apical branching of growing hyphae, the tip growth complex and the growing tip itself split in two. H: Simulation at elevated GTPase (*T*) and GAP (*T*_*g*_) levels (+) resulting in two clusters. I: Simulation starting in the one cluster regime (x) followed by a increase in *T* and *T*_*g*_ causing the single cluster to split. Time points (*t*) of snapshots are indicated inside each plot. All simulations domains have periodic boundary conditions in both directions. Colourbars indicate active GTPase concentrations ([GTPase]).

It has been suggested that lateral branching in fungi may be the result of apical dominance factors in the tip that suppress branching in the vicinity [44, 45]. However, our finding that turnover or negative feedback is needed to prevent polarisation (Fig 2) suggests that apical dominance may well be the default state and not require any dominance factor. Inhibitors (e.g. GAPs) may well be involved, but rather than merely creating an inhibition zone where no new clusters can be formed, their main role could be to keep the existing cluster from expanding indefinitely, thereby actually enabling the formation of new clusters. Alternatively, at large distances from the tip, it may no longer be reasonable to assume mass conservation across the entire hypha, and GTPase turnover will break competition, allowing a new tip to form.

Apical branching requires the splitting of an existing cluster rather than the appearance of a new one. Previously, an accumulation of inhibitor in the cluster has been suggested as potential mechanism [46]. Our results on the ODE model with GAP feedback suggest this might be possible, if we could significantly increase the amount of GAP without leaving the Turing regime. We can achieve this by simultaneously raising the total amount of GTPase (Fig 8G). At higher levels of both total GTPase and GAP compared to the single cluster set-up used before, we indeed find that two stable clusters form (Fig 8F-H, S5 Video). Upon increasing the total amounts of GTPase and GAP to this level starting from a single cluster steady state, the single cluster splits in two (Fig 8I, S5 Video). This suggests that apical branching may occur by accumulation of GTPase and GAP from fusing vesicles during tip growth.

These results and considerations demonstrate that a mechanism that allows for stable coexistence can offer elegant explanations for a range of phenomena in tip growing systems that could not well be explained with a polarisation mechanism. The specifics of individual systems remain a topic for future investigation.

## Discussion

In this study, we uncovered and investigated several mechanisms through which highly similar GTPase-based systems can generate different types of patterns. Polarisation is the invariable result of a mass conserved GTPase under positive feedback activation, because the stronger activation in larger GTPase clusters leads to a gradient and corresponding net flux of inactive GTPase from smaller clusters to larger ones (Fig 9A). Stable coexistence of multiple GTPase clusters can be achieved either by breaking mass conservation, or adding negative feedback through the activation of an inhibitor. In the first case, a constant supply of fresh GTPase across the membrane allows smaller clusters to grow in spite of the net flux to larger clusters (Fig 9B). In the latter case, larger clusters activate more inhibitor, limiting their growth (Fig 9C). In contrast to a previously proposed mechanism based on saturation of self-activation, these mechanisms lead to actually stable coexistence and can also explain the emergence of additional clusters as occurs, e.g., during branching in tip growth. Our use of two different minimal models suggests that these conclusions do not depend on the precise positive feedback mechanism.

**Fig 9.**
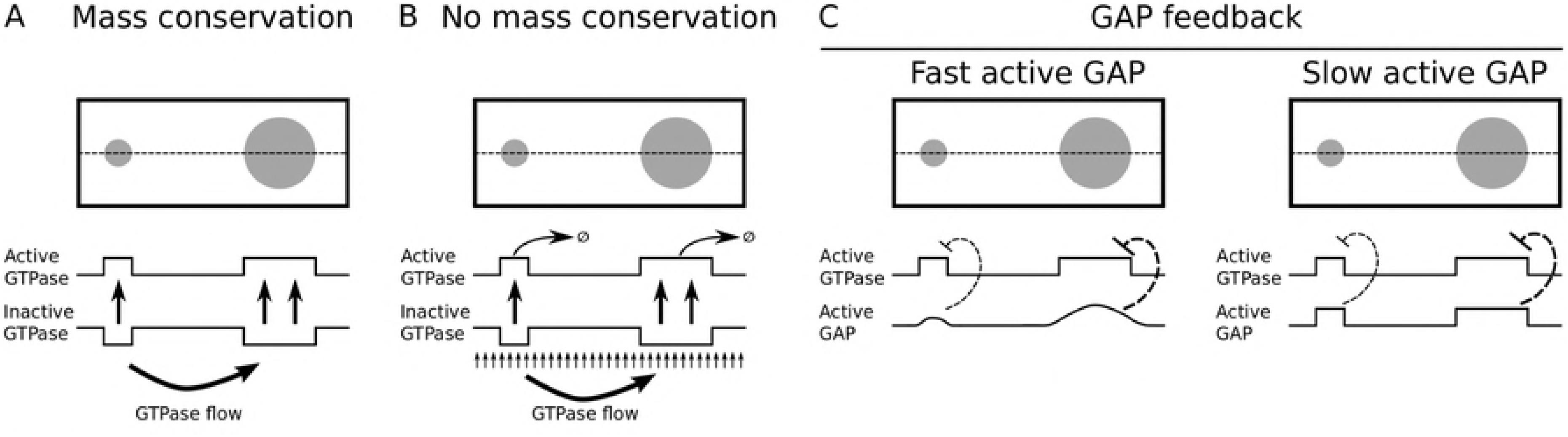
Proposed mechanisms for polarisation and coexistence. A: In polarisation models, larger clusters of active GTPase more effectively deplete the local supply of inactive GTPase than smaller ones. This results in a gradient of inactive GTPase that favours diffusion towards the larger cluster allowing the larger cluster to grow at the expense of the smaller one. B: When mass conservation is broken, production provides a fresh supply of GTPase to sustain smaller clusters in spite of this flux while degradation prevents larger clusters from growing uncontrollably, enabling coexistence. C: When active GTPase promotes GAP activation, larger clusters activate GAPs more effectively, promoting their own inactivation and allowing smaller clusters to survive. However, if GAPs were to diffuse as slowly as active GTPase, they would not be able to escape the cluster and quench the ability of the system to give rise to patterns in the first place.

### Mass conserved polarisation models cannot explain all relevant membrane patterns

Most existing models for polarisation involve mass conservation and some form of positive feedback [5]. Our results show that these two properties indeed consistently result in polarisation for both direct positive feedback and double negative feedback. Analysis of a simplified ODE approximation of a system of competing clusters suggests that this happens because larger clusters more effectively deplete their local reserve of inactive GTPase (Fig 9A). In the ODE model, the local pool of inactive GTPase is considered homogeneous. In reality, the level of inactive GTPase in the cluster will be lower than the level surrounding the cluster and inactive GTPase will flow towards both clusters from their direct surroundings. However, even in the worst case scenario the larger circumference of the larger cluster will cause more inactive GTPase to diffuse to the larger cluster. On a 1D domain, this effect would not be apparent. This insight may explain the longer competition times in 1D as compared to 2D.

A recent theoretical study proposed that severely slowed competition as a result of saturation could explain the difference between polarisation and coexistence [35]. On top of its inability to explain the appearance of new clusters after an initial pattern has been established, our findings show that this mechanism works less well when the formation of many clusters at the same time is considered. Therefore, the mechanisms for stable coexistence we propose here provide a better explanation for patterns as found in metaxylem [19], pavement cells [34], neurons [4], and fungal hyphae [27].

### Two types of model extensions can stabilise coexisting clusters

#### Sufficiently strong GTPase turnover can stabilise coexistence

The main difference between existing polarisation models that generate a single cluster and classical activator substrate depletion models that generate multiple clusters [12] is the assumption of mass conservation. We showed in 2D that by simply adding production and degradation terms to the two polarisation models, stable steady states with multiple clusters can be obtained as has been shown before for the 1D case [37]. Our ODE approximation suggests that this coexistence results from a redistribution of GTPase, with degradation mostly affecting larger clusters while production is homogeneous (Fig 9B). Moreover, we found that stable coexistence requires degradation of the *active* form. If only the inactive form is degraded, active GTPase in clusters can escape degradation and clusters keep growing until they fill the entire domain.

Broken mass conservation can only explain patterns of coexisting clusters if there is sufficient GTPase turnover for a given domain size. When the turnover rate or domain size becomes too small, models with turnover will converge to mass conserved models and polarisation will occur. This explains, for example, how a study on root hair initiation could obtain a single GTPase cluster with a model containing production and degradation terms [47]. To obtain stable coexistence, half-lives should most likely be shorter than the 10 to 30 hours reported for GTPases from a macrophage cell line [48]. However, different GTPases may have different turnover rates, and even half-lives of the same GTPase may be altered significantly by regulation [49], so this does not seem unreasonable.

Stable coexistence through GTPase turnover seems especially plausible on large domains, such as plant cells. In contrast, mass conservation is more plausible on smaller domains, such as neuron cell bodies [50], so that other mechanisms may be required to explain coexistence there. Since larger domains can hold more stable clusters, broken mass conservation may also be able to explain the formation of new clusters in between existing ones on a growing domain. This may explain the appearance of new protrusions during lateral branching of hyphae [43] and in growing pavement cells [34]. However, this behaviour does not provide the cell with much dynamic control, as it links cluster number to the domain size for any given turnover rate.

#### GAP feedback is a flexible alternative for stabilising coexistence

Even in cases of (near) mass conservation, coexistence is still possible through further additions to the interaction motif. Our results show that addition of negative feedback through activation of a sufficiently fast diffusing inhibitor (GAP) can stabilise coexistence in the two polarisation motifs studied. This is consistent with previous suggestions [38]. The ODE model indicates that GAP feedback fulfils this role by punishing larger clusters, which activate more GAP (Fig 9C). This mechanism only works when GAPs diffuse faster than active GTPase, possibly because too slowly diffusing GAPs will too strongly accumulate locally in clusters and extinguish them. This difference in diffusion rates could be achieved if active GAPs are not membrane-bound, or at least do not interact as strongly with the membrane or membrane-bound proteins as active GTPase.

GAP feedback and broken mass conservation are not mutually exclusive and which interaction motif is used in practice will have to be judged on a case by case basis. Experimental evidence suggests GAP feedback is involved in the spotted pattern found in metaxylem [19]. Our modelling results predict that if this feedback is indeed responsible for the coexistence of multiple GTPase clusters, experiments reducing GAP expression would result in fewer clusters.

Unlike broken mass conservation, GAPs provide extra options for regulation, making them more flexible. As shown by our tip splitting simulations, simultaneously providing extra GTPase and GAP can result in the splitting of a GTPase cluster. Previously, dilution due to repeated fusion of vesicles during tip growth has been suggested as a source of negative feedback to achieve tip splitting [46], but it would be hard to combine this with an increase in GTPase levels. Regulating GAP feedback, however, gives the cell the ability to control the number of clusters, even independent of the domain size.

Another example can be found in fission yeast (*Schizosaccharomyces pombe*), where a bipolar pattern of two active GTPase clusters promotes growth of the rod shaped cells in both directions. Upon cell division, both daughter cells start with a unipolar pattern and grow in a single direction until a certain size is reached at which a second GTPase cluster forms and bipolar growth is resumed (“new end take off”; NETO) [51]. The appearance of an additional cluster on a larger domain could be explained by both types of coexistence models. However, to also explain the reported oscillations between both tips requires a time delayed negative feedback [52, 53], which could not be achieved through linear GTPase degradation, whereas an extra molecular player (such as GAP) offers more flexibility to introduce a delay.

### Dynamic regulation of GTPase patterns in tip growing systems

Our findings suggest that multiple as yet poorly understood phenomena in mycelial tip growth could be explained by assuming some form of (GAP-like) negative feedback as is also implicated in pollen tube growth [28, 54]. Such negative feedback could, for instance, explain the occasional disappearance and reappearance of the tip growth complex observed in *A. gossypii* [29]. A combination of regulated negative feedback and an increase in total GTPase may also explain apical branching observed in this species [43]. Root hairs in plants do not normally branch, but overexpression mutants of ROP2, the GTPase controlling tip growth in root hairs, have root hairs with strong apical branching [55]. This supports the hypothesis that an increase in total GTPase, combined with some form of negative feedback, can result in apical branching through splitting of the GTPase cluster at the tip.

Since our models show that a single cluster is obtained unless sufficient turnover or GAP feedback is involved, polarisation may well be the default state. In this case, hypothesized apical dominance factors [45] that suppress branching would not be required. Rather, there would be more need for a branching signal that either stimulates negative feedback or GTPase turnover. Indeed, for arbuscular mycorrhizal fungi, a branching signal seems to be present in the form of strigolactones, although the precise molecular mechanism is still poorly understood [56]. Therefore, studies on hyphal branching focusing on identifying and characterising such branching factors may prove more fruitful than studies looking for apical dominance factors.

## Materials and methods

### Initial conditions

We initiated PDE simulations at the homogeneous steady state (see section 2 in S1 Appendix) with an amount of noise added to each integration pixel for the active form and the same amount subtracted from the corresponding inactive form. This made it as if a random small amount was interconverted between active and inactive form, without changing the total mass at each pixel. Per pixel, the noise was drawn from a normal distribution with a mean of 0 and a standard deviation of 10^*−*6^.

### Numerical methods

We performed numerical simulations using the python package Dedalus [57], which implements a spectral solver method, with the recommended dealias factor of 1.5 and the Runge-Kutta time-stepper. Fourier and Chebyshev basis functions were used for the x- and y-direction respectively, except for single cluster simulations, where Fourier basis functions were used in both directions. To determine appropriate temporal and spatial step sizes, we first performed several trial simulations for each model with reproducible perturbations as previously described [58], so that accuracy could be assessed using mesh refinement and time step reduction. We performed the final simulations with noise added directly to each integration pixel to ensure all possible wave lengths are represented. Integration steps used for final simulations are given in S1 Table. We continued all simulations until a steady state was reached (no more noticeable changes in the concentrations). In some simulations, a stable pattern ended up drifting at a constant speed in the periodic direction. This can happen because with periodic boundary conditions any shift of a solution is also a solution. Therefore, such drifting patterns were regarded as steady states.

We performed simulations of the ODE models in matlab using the function ode45 with default parameters.

### Bifurcation and stability analysis

For the models with only two states (WP and WPT), we performed both a classical linear stability analysis (LSA) and the asymptotic local perturbation analysis (LPA). For the remaining models, LSA is not feasible and only LPA was used. LSA can be used to determine under what conditions arbitrarily small spatial perturbations in a homogeneous state can grow. This way, parameter regimes where spontaneous patterning occurs can be identified. The wave numbers of the perturbations that become unstable have often been used to predict the length scales of the pattern, but these are only valid close to the homogeneous state and therefore not in general a good reflection of the length scales of the final pattern [39]. We performed LSA as previously described [59] as described in section 3 of S1 Appendix.

LPA is a recently developed asymptotic analysis for reaction-diffusion models [60, 61]. It works by considering the behaviour of a local pulse in the activator concentration, in the limiting case where the diffusion coefficients of slowly diffusing components approach zero and those of rapidly diffusing components approach infinity. This reduces the system of PDEs to a system of ODEs that can be analysed with existing bifurcation software. It is, therefore, not as exact as LSA, but it can be more easily scaled up to models with more than two components and it can also be used to chart the areas of parameter space where the homogeneous state is stable, but coexists with a stable heterogeneous state. In our case, we used strong differences (100 fold) in diffusion rates and, therefore, regimes predicted by LSA and LPA matched quite closely. We performed LPA on all our models as described by others [60] (see section 4 of S1 Appendix for details) and analysed the resulting ODEs using the matcont package for matlab [62].

### Single cluster simulations

To study phenomena observed during tip growth, we performed simulations with the same parameters as before, but on a smaller domain, such that only a single cluster formed. For these simulations we used a square domain with periodic boundary conditions on all sides. This domain represents the tip of the growing tube. The dimensionless domain size was 19.0 × 19.0 for the WPGAP models, and 31.6 × 31.6 for the WPT model. To ensure that any unstable states reached would be disrupted, we added noise not only at the beginning, but also every 10 time units. This noise was also drawn from a normal distribution with a mean of 0 and a standard deviation of 10^*−*6^.

## Supporting information

**S1 Fig. Model simulations with concentrations and time points.**

Simulations are as in Fig 2, but concentration ranges and time points at which simulations were stopped are indicated.

**S2 Fig. Linear stability analysis for models with two variables.** When the real part of at least one eigenvalue (*λ*) corresponding to a certain admissible wave number (*k*) is greater than zero the homogeneous state is unstable and a pattern forms. Admissible wave numbers for the geometry of the simulations are indicated with red crosses (top figures) for various parameter values. Green lines show the maximum real part of *λ* as a function of total GTPase (*T*, WP model) or GTPase production rate (*σ*, WPT model), plotted both against these parameters (bottom) and against the squared wave numbers (top). Cyan lines indicate real parts of complex eigenvalues where present.

**S3 Fig. Reduced turnover decreases the number of coexisting clusters generated by the WPT model.** Steady state (*t* = 200000) active GTPase profiles generated by the WPT model with production (*σ*) and degradation (*ξ*) rates reduced by a factor 10, 100, and 1000 compared to default parameters.

**S4 Fig. Two parameter bifurcation plot for the WPGAP model with low active GAP diffusion from Fig 5C.** Crosses indicate parameter settings where trial simulations were performed. All simulations resulted in polarisation.

**S5 Fig. Tip growth simulations with the WPT model.** B: Bifurcation plot with symbols showing parameter settings used in the simulations, corresponding to symbols over simulations and cartoons. For meaning of lines see Fig 5. A: In pollen tubes, growth occurs in pulses as negative feedback during growth results in an oscillatory GTPase cluster at the growing tip. C: Simulation of a single active GTPase cluster with increase in *ξ* upon reaching steady state followed by a return to starting levels. D: In growing hyphae of *A. gossypii*, the tip growth complex sometimes disappears corresponding to a halt in growth, suggesting involvement of negative feedback. E: Simulation with two subsequent increases in *ξ*, followed by a return to the starting level. Symbols over simulations correspond to those in the bifurcation plot. Time points (*t*) of snapshots are indicated inside each plot. All simulations domains have periodic boundary conditions in both directions. Colourbars indicate active GTPase concentrations ([GTPase]).

**S1 Table. Spatial and temporal integration step sizes used for numerical integration.**

**S1 Appendix. Supplementary text.** Including non-dimensionalisation, derivation of homogeneous steady states, details of LSA and LPA methods, analysis of the WPT model with degradation of active GTPase, detailed derivation of the ODE models, and the quasi steady state approximation used for the ODE model with GAP feedback.

**S1 Video. Model simulations.** Time lapse movies of model simulations from Fig 2, showing concentrations of active GTPase. For models with two GTPases only concentrations of the first are shown. The concentration profile of the second GTPase is always complementary to that of the first.

**S2 Video. Model simulations in Hopf regimes.** Time lapse movies of model simulations in Hopf regimes for the WPT and WPGAP models. Concentrations of active GTPase are shown. The WPT simulation was performed with parameters *σ* = 0.4 and *ξ* = 1, domain height *H* = 316, and domain width *W* = 190. The WPGAP simulation was performed with parameters *T* = 34.83 and *T*_*g*_ = 100, domain height *H* = 50 and domain width *W* = 30. All other parameters were at default values.

**S3 Video. Simulation WPT model with degradation of inactive GTPase.** Time lapse movie of model simulation described in section 5 of S1 Appendix, showing concentrations of active GTPase.

**S4 Video. Simulation WPT model with degradation of both active and inactive GTPase.** Time lapse movie of model simulation described in section 7 of S1 Appendix, showing concentrations of active GTPase.

**S5 Video. Simulations of tip growth scenarios.** Time lapse movies of model simulations from Fig 8 and S5 Fig, showing concentrations of active GTPase.

**S1 Code. Scripts used to generate the figures.**

## Acknowledgments

We thank Marcel Janson for useful comments on the tip growth case study.

